# Experience-dependent increases in sharp-wave ripples near visual targets

**DOI:** 10.1101/054171

**Authors:** Timothy K. Leonard, Kari L. Hoffman

**Affiliations:** Department of Psychology, York University, Toronto, ON, Canada; Neuroscience Graduate Diploma Program, York University, Toronto, ON, Canada; Centre for Vision Research, York University, Toronto, ON, Canada; Department of Biology, York University, Toronto, ON, Canada

## Abstract

We measured hippocampal sharp-wave ripples during goal-directed visual exploration in macaques. Exploratory sharp-wave ripples were more frequent on familiar trials, in the second half of search, and near the visual target (i.e., the goal location). These spatial and temporal properties may help SWRs coordinate hippocampal and extra-hippocampal firing sequences that guide actions based on past experiences.

---

Hippocampal sharp-wave ripples (SWRs) are nested high-frequency oscillations that occur in a range of mammalian species during slow-wave sleep, quiet wakefulness and brief pauses in exploration (Girardeau & Zugaro, 2011; Buzsáki, 2015). In rodents, the production of SWRs aids in memory consolidation during quiescence (Girardeau, Benchenane, Wiener, Buzsáki & Zugaro, 2009; Ego-Stengel & Wilson, 2010; Dupret, O’Neill, Pleydell-Bouverie & Csicsvari, 2010). During waking behaviors, SWRs support spatial coding and memory expression (O’Neill, Pleydell-Bouverie, Dupret & Csicsvari, 2010; Carr, Jadhav & Frank, 2011; Jadhav, Kemere, German & Frank, 2012; Pfeiffer & Foster, 2013; Singer, Carr, Karlsson & Frank, 2013). Specifically, the occurrence of SWRs at goal locations predicts rats’ memory performance (Dupret, O’Neill, Pleydell-Bouverie & Csicsvari, 2010). In primates, SWRs additionally occur during active, goal-directed visual search (Leonard et al., 2015), though the role of these (or any) hippocampal SWRs in primate memory has not been established. We therefore recorded SWRs from the macaque during a memory-guided visual search task, to determine whether or not exploratory SWRs varied with learning, and as a function of search distance to the goal (target) location.

Macaques searched for target objects embedded in scenes (Figure 1a, b), revealing faster detection – speeded search – with scene repetition (Figure 1c; Z(1120) = −13.439, P < 4.844e-41, online methods), indicating relational/binding memory for items-in-context (Chau, Murphy, Rosenbaum, Ryan & Hoffman, 2011). Hippocampal recordings taken during task performance showed that exploratory sharp-wave ripples (eSWRs) occur at a higher rate on repeated trials than novel trials (figure 2a; P <= 0.0001, permutation test). To see when this SWR-enhancement emerges in the trial, we calculated ripple rates from scene onset separately for novel and repeated presentations (Figure 2b). Repeated trials had higher ripple rates beginning shortly into the trial, and were not restricted to narrow temporal windows. After accounting for differences in trial duration associated with successful trials and repeated trials, there was still an overall tendency for eSWRs to appear in the final half of search ‘journeys’ rather than in the initial half (Table 2, online methods).

**Figure 1.**
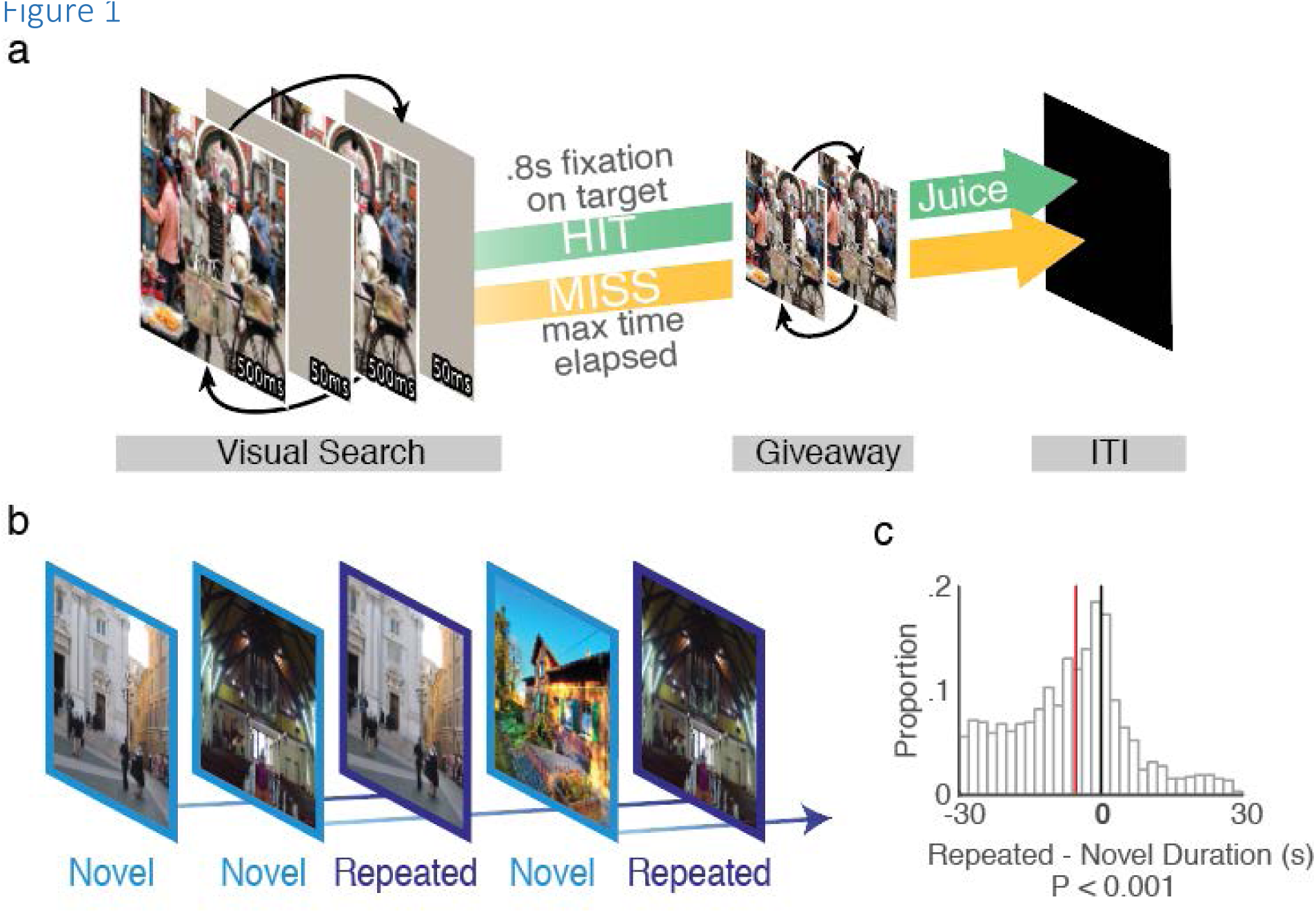
Memory-guided visual search task. (**a**) Trial structure. Left: an original and target-modified scene are presented in alternation, with a brief mask presented in between, requiring protracted visual search on the image to detect the changing target (’flicker change detection’). Visual search ends after the target is fixated for .8 seconds or the maximum on-screen search time is reached. Following visual search is the ‘giveaway’ (middle), consisting of an alternation between the two scenes, without mask, revealing the changed location, and ending with an inter trial interval (ITI) black screen period (right). (See online methods, Hoffman et al., 2013, Leonard et al., 2015, for more details). (**b**) A sequence of flicker scene pairs are presented in blocks of 30 trials. Novel-trial scenes were never presented to that animal before, and these comprised 12/30 trials per block. Each novel-trial scene was repeated after at least 1 but not more than 36 intervening trials, such that repetitions occurred either within a single block or in the next trial block. (**c**) Histogram of trial duration changes with scene repetition. Targets are typically found faster on repeated trials of a scene than on novel presentations (i.e. trial durations are shorter) indicated by the leftward, negative bias; red line indicates the median difference value (−5.27 s).

**Table 2.** **Table 2**Logistic regression results, predictors of SWRs occurring in 1^st^ or 2^nd^ half of trials.

| Coefficients | Estimate | SE | T | P |
| --- | --- | --- | --- | --- |
| (Intercept) | -2.9314 | 0.9088 | -3.2254 | .005* |
| Trial Success | 0.7132 | 0.4892 | 1.4580 | .193 |
| Trial Type | 0.2012 | 0.3806 | 0.5287 | .597 |
| Trial Duration | 0.0728 | 0.0239 | 3.0480 | .005* |

Note: \* $P < 0.01$ ; Trial Success: Factor, Miss (id 0; n: 77), Hit (id 1; n: 68); Trial Type: Factor, Novel (id 0; n: 49), Repeated (id 1; n: 96); Trial Duration: seconds; 145 observations, 141 error degrees of freedom; $\chi^2$ : 11, $P = 0.0115$

SWR occurrence near goal locations has been associated with learning (Dupret, O’Neill, Pleydell-Bouverie & Csicsvari, 2010), thus eSWRs during repeated-scene search may be more frequent when gaze is focused near the target. To test this, we measured the distance between a scene’s target object and the gaze position (’fixation location’) at the time of an SWR, grouped according to novel/repeated trial type. Because the prior distribution of fixation distances to targets may not be flat, and could vary with trial repetition, we compared the SWR-locked distances of a given trial type to *all* fixation-to-target distances of that trial type. SWR-locked fixation distances to the target are closer on repeated than novel trials (Figure 3a), even after considering the prior distribution (non-SWR-locked fixation distances with repetition; two way ANOVA on fixation-to-target distances; trial type main effect: P < 0.0011; SWR main effect P = 0; trial type x SWR interaction: P = 0.001; see Table 1; baseline (non-SWR)-normalized distances for each condition, Figure 3b; Z(266, 330) = 3.2604, P = 0.0011).

**Figure 2.**
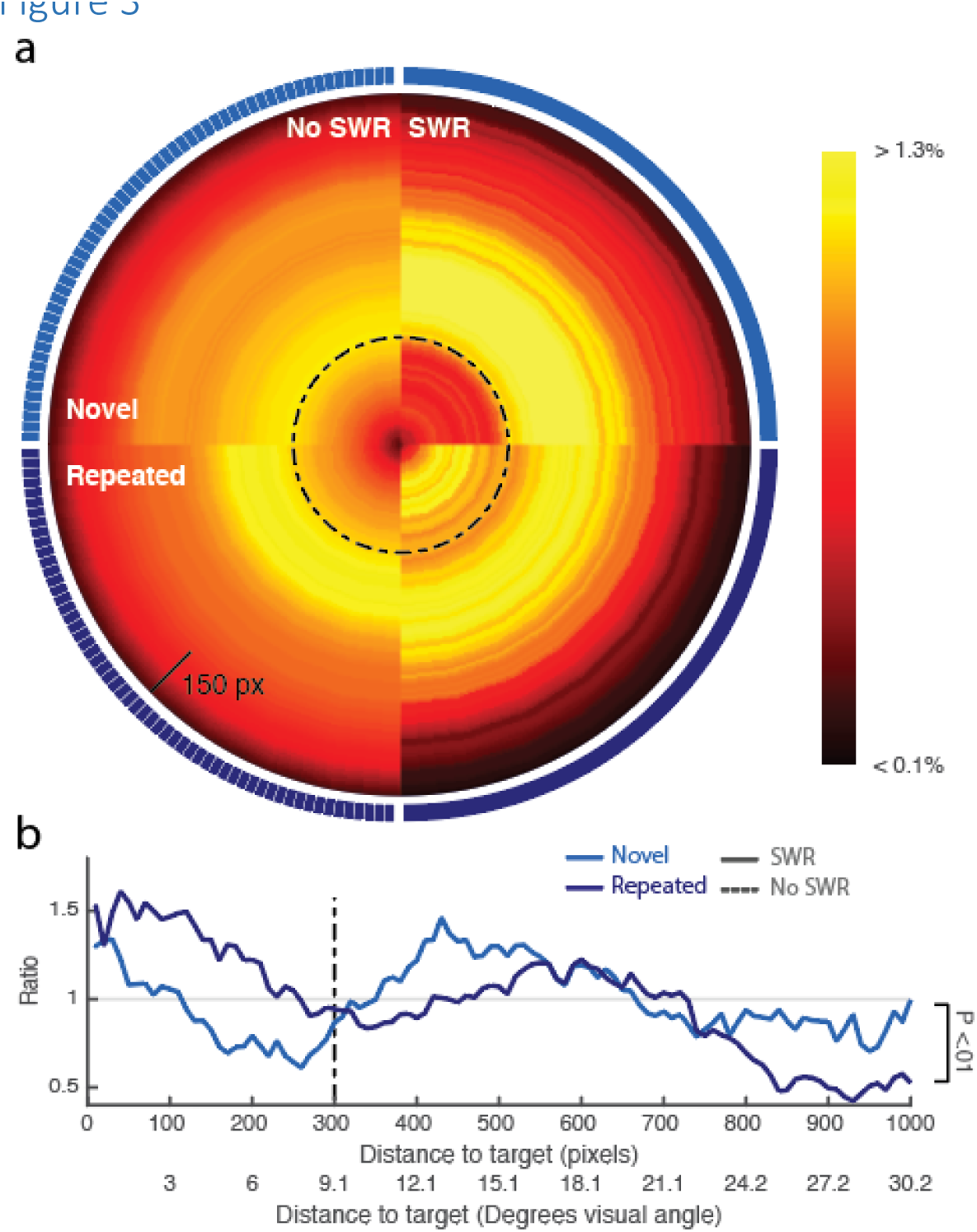
SWR occurrence increases near the target on repeated trials. (**a**) Distance of fixations from the target, centered at 0 pixels. Upper left quadrant: all novel-trial fixations; Upper right quadrant: novel-trial fixations locked to SWRs; Lower left quadrant: all repeated-trial fixations; Lower right quadrant: repeated-trial fixations locked to SWRs. Black dotted line indicates a distance of ~9 degrees of visual angle (300 pixels) from the nearest target edge. Color values represent percentage of fixations, for given condition, at that distance from target edge (see online methods). (**b**) Frequency of SWR-locked fixation distance to target, normalized by all fixation distances to target, of a given trial type (Novel, Repeated). 0 indicates the edge of the target object; the dotted line at 300 pixels matches the dotted line in **a**. Trial type, SWR occurrence, and their interaction, were all significant predictors of distance of fixation to target edge (Two-way ANOVA; see table 1 and online methods).

**Table 1.** **Table 1**ANOVA results, Fixation distance to Target by trial type and during/without SWR. See online methods for post hoc validation of P values.

| Trial Type | SWR | Mean | Median | SD | n |
| --- | --- | --- | --- | --- | --- |
| Novel | No SWR | 525.01 | 501.11 | 292.66 | 12631 |
| Novel | SWR | 510.74 | 488.38 | 274.61 | 266 |
| Repeated | No SWR | 525.41 | 494.14 | 293.05 | 11800 |
| Repeated | SWR | 430.95 | 428.23 | 242.28 | 330 |
| Source | Sum Sq. | d.f. | Mean Sq. | F | P |
| Trial Type | 906649.7 | 1 | 906649.7 | 10.63 | .0011* |
| SWR | 1700374.5 | 1 | 1700375 | 19.94 | 0** |
| Trial type x SWR | 924776.8 | 1 | 924776.8 | 10.84 | .001* |
| Error | 2134335235 | 25023 | 85294.9 |  |  |

Note: \* $P < 0.01$ , \*\* $P < 0.001$ ; Table values (top) are expressed as distance in pixels from fixation to target edge.

In rats, goals are over-represented in both the frequency of SWRs and in ripple-associated internally-generated sequences, even when initiated remotely from the goal (Pfeiffer & Foster, 2013). Furthermore, receptor blockade suggests this is due to NMDA-dependent plasticity during learning (Dupret, O’Neill, Pleydell-Bouverie & Csicsvari, 2010). During sequence replay of location, specific cell firing can be biased by current in-field cells (Diba & Buzsáki, 2007; Pfeiffer & Foster, 2013). Although speculative, such a bias in the present study could increase gaze-location-selective firing near the target, facilitating later recall of the target location. In addition to influencing local ensembles, SWRs can have widespread effects on neocortical and subcortical structures (Logothetis et al., 2012) while targeting specific subsets of cells in specific target areas (Ciocchi, Passecker, Malagon-Vina, Mikus & Klausberger, 2015). Thus, the role of SWRs in waking tasks may relate to memory retrieval and generation of goal trajectories (Roumis & Frank, 2015) affecting extra-hippocampal activity (Jadhav, Rothschild, Roumis & Frank, 2016). Our results show that exploratory SWRs observed in primates (Leonard et al., 2015), are enhanced during learning, specifically near visual targets, and thus the SWR-mediated mechanisms that support route finding and spatial decision-making in rodents may be co-opted in primates to support non-ambulatory visual, and episodic-like learning.

## Online Methods

### Electrophysiology

All procedures were conducted with approval from the local ethics and animal care authorities (Canadian Council on Animal Care). These methods are described in detail in Leonard et al., (2015); M1 and M2. Animals (2 adult female rhesus macaques; *M. mulatta*) were implanted with platinum/tungsten multicore tetrodes (Thomas Recordings, Giessen, Germany), lowered into the CA3/DG region of hippocampus. Location was based on MR images (M1), on MR/CT coregistration of electrode position (M2), and on functional characterization based on expected white/gray matter and ventricle transitions in depth during lowering. In addition, we observed SWRs only in limited ranges of depth, within which we encountered the best single unit isolation.

Neural recordings for field potentials were digitally sampled at 32 kHz using a Digital Lynx acquisition system (Neuralynx, Inc., Bozeman, Montana, USA), filtered between 0.5 Hz and 2 kHz.

SWR events were detected by an observer selecting the tetrode and channel with the clearest ripple activity, filtering that LFP signal (100-250 Hz), transforming to z-scores, rectifying, and band pass filtering 1-20 Hz. Events which registered 3 SD above the mean, with a minimum duration of 50ms beginning and ending at 1 SD, were identified as ripples.

### Task

The task, performed by both subjects, consisted of flicker change detection of an object in a natural scene, previously shown to be a hippocampal-dependent relational memory task in macaques (Chau, Murphy, Rosenbaum, Ryan & Hoffman, 2011). The flicker change detection task consisted of an original scene alternated with an object-manipulated scene, with a short mask (grey screen) shown between each image presentation, making the changing object difficult to identify without prior experience with that flicker scene. Visual search on the flickering scene lasted until the changing object was fixated for at least .8 s (target found; ‘hit’) or a total search time maximum was reached (target not found; miss). Maxima were 45s for M1, and 30s for M2, which allowed faster progression through the sets in M2. Analysis was restricted to 30-s trials of M1, for comparison across animals. Found targets were rewarded with a preferred juice reward, and all trials entered a black-screen inter-trial interval before the next trial started (ITI: 20s, M1; 8s, M2). If the subject looked off screen the trial was paused. If the amount of off-screen looking exceeded the total trial duration (all off-screen and on-screen looking) by 50% that trial was removed. Previous work demonstrated that rapid repeated-trial search corresponded to explicit memory for unique object-scene pairs in humans, and that macaques reliably show rapid search with repetition (Chau, Murphy, Rosenbaum, Ryan & Hoffman, 2011).

### Statistics

#### Novel-Repeated paired search durations

Novel image trials were matched to their repeated image counterparts from all sessions with SWR events (M1 (LE): 36 daily sessions, M2 (LU): 45 daily sessions). Where an image was repeated twice, each repetition is paired with the novel trial duration. This resulted in 1218 novel/repeated trial pairs consisting of 591 novel presentations with 761 counterparts. From this collection of 1218 pairs, 98 pairs, all from M1 (LE), were excluded due to having at least one trial of pair being over 30 seconds in duration. Repeated trial durations (mean 15.163, std 10.256) were subtracted from the trial duration for the corresponding novel images (mean 9.045, std 8.834). Because these difference values were non-normally distributed (One sample KS test; KS(1218) = 0.5859, P < 0.001, two-tail), a non-parametric paired-samples test (Wilcoxon signed-rank test, two tailed) was used to test if the distribution of repeated-novel durations were different than 0.

#### Novel-Repeated Rate comparisons

Rates for 584 novel and 749 repeated trials, with 49 and 96 SWRs, respectively, where both novel and repeated trial durations were under 30 seconds, were compared using a label-swap permutation test, with a two-tail design. Rates are calculated from trial durations and numbers of SWRs per condition. To statistically test differences between these conditions, we compared the observed mean rate difference between novel and repeated conditions (mean SWR/min: .306; mean SWR/min: .708), to 10000 mean rate differences generated by randomly swapping condition labels for the trial duration, and SWR occurrence, data.

##### Rates over time

For all periods of on screen search behavior (all off-screen looking behavior removed), the SWR rate was calculated using a sliding window (width 4s, steps 1s; 0 to 30s; 27 windows for each condition). The rate was calculated, in a given window, as the number of SWR events divided by all trial durations which fell into that window.

Differences in rates were tested using a permutation test, comparing the differences distribution for Novel vs. Repeated rates for each sliding window. To statistically test differences in rates between conditions, tests were run with 10,000 permutations, for each window, establishing the probability of the observed difference in novel vs. repeated rate having occurred, using a two tail design. These permutation test values were then corrected using the positive false discovery rate method (pFDR; Storey, 2002).

Figure 2b shows variability estimated for each window of each condition with the bootstrap method (sampling with replacement), whereas the permutation test statistic was the mean difference between conditions, generated with a label swap procedure (sampling without replacement). Because the significance test measures mean differences between conditions, and the confidence intervals reflect variability within conditions, the 95% bootstrap confidence intervals may overlap where permutation test shows significance.

**Figure 2.**
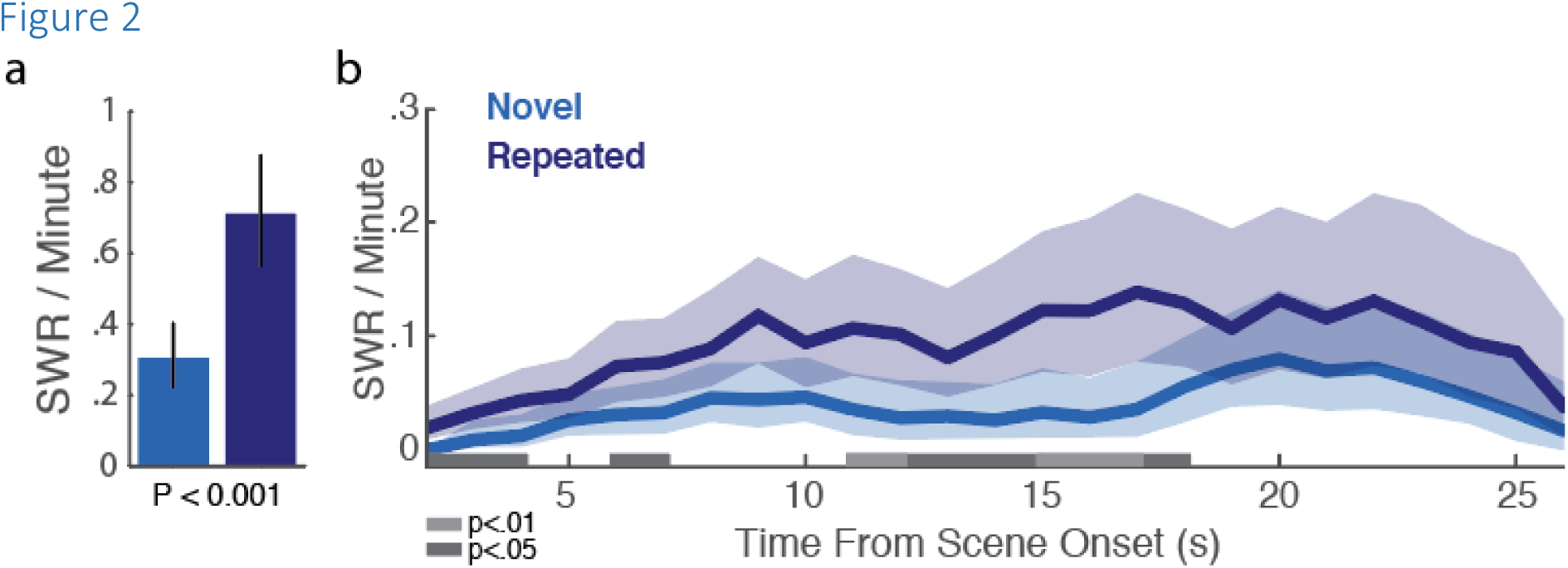
SWR rates increase with trial repetition. (**a**) Mean SWR rates for novel (blue) and repeated (purple) trials. Error bars are 95% bootstrap confidence intervals (n=1000; P<0.001). (**b**) SWR rates over time, for Novel and Repeated trials. Shaded portions represent 95% bootstrap confidence intervals (n=1000). Periods in which rates differ, after surviving multiple comparison correction, are highlighted in gray (see methods).

#### Two way ANOVA

556 fixations, which occurred within +- 250ms of the center of an SWR event were labelled as fixations with SWR occurrence, 24431 fixations which occurred 5s before or after an SWR event were labelled as fixations without SWR occurrence. For all fixation events, the distance from the fixation to the closest edge of the target bounding box was calculated. A two way ANOVA compared the effect of trial type (Novel/Repeated), occurrence of SWR and interaction between SWR and trial type on the distance of the fixation to target edge. See table 1.

Levine’s test showed that at least one of the groups had a significantly different variance (P<0.001), and so further validation of the ANOVA results was performed.

##### ANOVA validation

Because the ANOVA is especially vulnerable to uneven variance when the groups are not equal size, we ran a permutation test to validate the ANOVA results. 10,000 label swaps were performed, preserving the unequal group sizes (see Table 1), and F values were only calculated for label swaps in which Levine’s test showed a significant difference in variability (p<0.001) between groups. The magnitude of the originally observed F values were then compared to this distribution of generated F values, where variability between groups was always met.

This validation resulted in P values (probability of actual observation having occurred), based on the observed F values compared with the generated distribution of F values, are within range of the parametric ANOVA results reported in Table 1. (trial type: p=.0045, SWR: p=0, trial type x SWR: .0034). The uneven sample sizes did not play role in inflating the effect sizes.

##### Post Hoc Tests

To test the interaction, where Repeated eSWR fixation locations are closer to the target than novel eSWR fixation locations, we compared distances from fixation to target, during eSWR events, for each trial type, baseline normalized by that trial type’s fixations *without* eSWR, and compared the distributions with a Wilcoxon Ranksum test. The Wilcoxon Ranksum (non-parametric test) was required as both normalized distributions were non-normal (Novel, one-sample KS test; KS(330)= 0.526, P = 6.1227e-86; Repeated, one-sample KS test; KS(266)= 0.516, P = 1.0333e-62).

#### Logistic Regression

For each trial with SWRs (145 observations) an index was calculated by dividing the time of the last SWR of a trial by the total trial duration. This index, representing time from SWR to end of trial, with range 0-1, describes SWRs near end of trial with low values approaching 0, and SWRs near start of trial with values approaching 1.

This index was then converted to dichotomous value, where values < 0.5 (those occurring before halfway through the trial) are labelled early SWR, and values >= .5 (those occurring beyond halfway through the trial) are labelled late SWR. The early or late occurrence of SWR was then predicted, using factors of Trial Success, Novelty and the continuous measure of Trial Duration with a logistic regression model. The logistic regression model was preferred over a linear one, due the underlying data lacking homoscedasticity.

The pairwise comparisons in table 2 are two tailed, and have been FDR corrected, using the Benjamini and Hochberg method (Benjamini & Hochberg, 1995).

## ACKNOWLEDGMENTS

This work was supported by the Krembil Foundation, Brain Canada, an NSERC Discovery grant, NSERC CREATE VSA and a QEII-GSST scholarship to TKL from the Ontario Ministry of Training Colleges and Universities.

## References

Benjamini, Y., & Hochberg, Y. (1995). Controlling the False Discovery Rate: A Practical and Powerful Approach to Multiple Testing. Journal of the Royal Statistical Society. Series B (Methodological), 57(1), 289–300.

Buzsáki, G. (2015). Hippocampal sharp wave-ripple: A cognitive biomarker for episodic memory and planning. Hippocampus, 25(10), 1073–1188.

Carr, M. F., Jadhav, S. P., & Frank, L. M. (2011). Hippocampal replay in the awake state: a potential substrate for memory consolidation and retrieval. Nat Neurosci, 14(2), 147–153. doi:10.1038/nn.2732

Chau, V. L., Murphy, E. F., Rosenbaum, R. S., Ryan, J. D., & Hoffman, K. L. (2011). A Flicker Change Detection Task Reveals Object-in-Scene Memory Across Species. Front Behav Neurosci, 5, 58. doi:10.3389/fnbeh.2011.00058

Ciocchi, S., Passecker, J., Malagon-Vina, H., Mikus, N., & Klausberger, T. (2015). Brain computation. Selective information routing by ventral hippocampal CA1 projection neurons. Science, 348(6234), 560–563. doi:10.1126/science.aaa3245

Diba, K., & Buzsáki, G. (2007). Forward and reverse hippocampal place-cell sequences during ripples. Nat Neurosci, 10(10), 1241–1242. doi:10.1038/nn1961

Dupret, D., O’Neill, J., Pleydell-Bouverie, B., & Csicsvari, J. (2010). The reorganization and reactivation of hippocampal maps predict spatial memory performance. Nat Neurosci, 13(8), 995–1002. doi:10.1038/nn.2599

Ego-Stengel, V., & Wilson, M.A. (2010). Disruption of ripple-associated hippocampal activity during rest impairs spatial learning in the rat. Hippocampus, 20(1), 1–10. doi:10.1002/hipo.20707

Girardeau, G., Benchenane, K., Wiener, S. I., Buzsáki, G., & Zugaro, M.B. (2009). Selective suppression of hippocampal ripples impairs spatial memory. Nat Neurosci, 12(10), 1222–1223. doi:10.1038/nn.2384

Girardeau, G., & Zugaro, M. (2011). Hippocampal ripples and memory consolidation. Curr Opin Neurobiol, 21(3), 452–459. doi:10.1016/j.conb.2011.02.005

Hoffman, K. L., Dragan, M. C., Leonard, T. K., Micheli, C., Montefusco-Siegmund, R., & Valiante, T. A., et al. (2013). Saccades during visual exploration align hippocampal 3-8 Hz rhythms in human and non-human primates. Frontiers in systems neuroscience, 7, 43. doi:10.3389/fnsys.2013.00043

Jadhav, S. P., Kemere, C., German, P. W., & Frank, L.M. (2012). Awake hippocampal sharp-wave ripples support spatial memory. Science, 336(6087), 1454–1458. doi:10.1126/science.1217230

Jadhav, S. P., Rothschild, G., Roumis, D. K., & Frank, L.M. (2016). Coordinated Excitation and Inhibition of Prefrontal Ensembles during Awake Hippocampal Sharp-Wave Ripple Events. Neuron, 90(1), 113–127.

Leonard, T. K., Mikkila, J. M., Eskandar, E. N., Gerrard, J. L., Kaping, D., & Patel, S. R., et al. (2015). Sharp Wave Ripples during Visual Exploration in the Primate Hippocampus. J Neurosci, 35(44), 14771–14782. doi:10.1523/JNEUROSCI.0864-15.2015

Logothetis, N. K., Eschenko, O., Murayama, Y., Augath, M., Steudel, T., & Evrard, H. C., et al. (2012). Hippocampal-cortical interaction during periods of subcortical silence. Nature, 491(7425), 547–553. doi:10.1038/nature11618

O’Neill, J., Pleydell-Bouverie, B., Dupret, D., & Csicsvari, J. (2010). Play it again: reactivation of waking experience and memory. Trends Neurosci, 33(5), 220–229. doi:10.1016/j.tins.2010.01.006

Pfeiffer, B. E., & Foster, D. J. (2013). Hippocampal place-cell sequences depict future paths to remembered goals. Nature, 497(7447), 74–79. doi:10.1038/nature12112

Roumis, D. K., & Frank, L.M. (2015). Hippocampal sharp-wave ripples in waking and sleeping states. Curr Opin Neurobiol, 35(Complete), 6–12.

Singer, A. C., Carr, M. F., Karlsson, M. P., & Frank, L.M. (2013). Hippocampal SWR activity predicts correct decisions during the initial learning of an alternation task. Neuron, 77(6), 1163–1173. doi:10.1016/j.neuron.2013.01.027

Storey, J.D. (2002). A direct approach to false discovery rates. Journal Of The Royal Statistical Society Series B, 64(3), 479–498.

